# Neuronal adaptation in the course of the prolonged task improves visual stimuli processing

**DOI:** 10.1101/2020.04.07.029959

**Authors:** Vladimir Maksimenko, Alexander Kuc, Nikita S. Frolov, Alexander Hramov, Alexander Pisarchik, Mikhail Lebedev

## Abstract

Brain optimally utilizes resources to resist mental fatigue during the prolonged period of cognitive activity. Neural mechanisms underlying long-term cognitive performance remain unknown. We show that during the 40-minutes visual stimuli classification task, subjects improve behavioral performance in terms of response time and correctness. We observe that the prestimulus *θ* and *α* power grows during the experiment manifesting the mental fatigue. The prestimulus *β* power, in its turn, increases locally in the region, engaged in the ongoing stimulus processing, that may reflect the neuronal adaptation. Our results evidence that the neuronal adaptation is enhanced in the course of the experiment reducing the cognitive demands required to activate the stimulus-related brain regions.

## Introduction

Since the brain resource is limited, human exhibits mental fatigue during the prolonged periods of cognitive activity. The mental fatigue negatively correlates with human attention and results in the decrease of the behavioral performance [1]. The bulk of literature describes the cortical activity underlying mental fatigue and attention. Thus, the mental fatigue is accompanied with the shift of EEG power toward the low-frequency bands, e.g. *δ* (1 − 4 Hz), *θ* (4 − 8 Hz), and *α* (8 − 13 Hz) [2, 3], while high-frequency activity, e.g. *β* (15 − 30 Hz) and *γ* (> 30 Hz) typically decreases in amplitude [4]. Attention, in turn, is characterized by the low *α*- and high *β*-band spectral power. In particular, low *α*- and high *β*-band power during the prestimulus period reflects increased attention and predicts better performance in the ongoing task [5, 6].

At the same time, cognitive performance is not necessary to decrease while accomplishing prolonged resource-demanding tasks. For instance, the increased reward makes the subject to improve behavioral performance even after the fatigue-induced session [7, 8]. It evidences that the brain spends cognitive resources in the way to reserve it for future use [9]. The Ref. [10] relates the strategy the brain utilizes the cognitive resources with a fatigue-induced performance decrement, but not with the subjective fatigue. It manifests that the resource reallocation is controlled by the endogenous cognitive mechanisms regardless of the subjective feelings. According to the recent review [11], the neural activity underlying the cognitive resource rearrangement and cognitive performance is yet to be determined.

In this work, we subject the group of volunteers to the prolonged (∼ 40 min) task. The task requires the participants to quickly percept the successively presented bistable visual stimuli and report one of their possible interpretations right after the presentation. On the behavioral level, subjects improve their performance in the course of the task: they reduce their response time along with the number of errors. We suppose that this observation can be referred to as the training effect and suggest that within-experiment training is one of the possible mechanisms of the optimal cognitive resource utilization.

The previous works reported that the training in the particular task allowed the subjects to improve their behavioral performance. In the visual task, training improved the efficiency of high-level visual processing, which therefore provided less ambiguous sensory information to the decision-related brain networks [12]. While in Ref. [12] the training period has lasted for three days; our results suggest that the training effect is notable even within a 40-min session.

Tang and Posner [13] previously described the brain’s ability to enhance its cognitive performance in the course of training. They distinguished two different types of training - network training and state training [14]. The network training involves the practice of a specific task (e.g., attention, working memory) and thus exercises the specific task-related brain areas and networks [15]. The state training involves exercises, e.g., meditation that might alter the brain state in general [16]. The network training, in its turn, may use a fixed task difficulty where performance shows improvement over trials or the adaptive training in which task difficulty adjusts as learning occurs [14]. Given the above, we suppose that our current study deals with the brain network training performed in the fixed difficulty fashion.

According to [14], the neuronal mechanisms underlying the training effect remain poorly understood. Some studies report the increased activation of the task-related brain networks in the course of training. In contrast, others observe the decrease of the task-related activity of the corresponding brain structures [17]. It also appears that the direction of the effect depends heavily on the type of training. For instance, the increased task-related brain activity may reflect the need for increased effort to solve more complicated tasks during adaptive training [18]. Thus, there is no complete understanding of how brain activity changes during the training.

To address this issue, we consider the cortical activity on the EEG sensor level in *θ* (4-8 Hz), *α* (8-12 Hz), and *β* (15-30 Hz) frequency bands in the course of the experiment. We observe that the training changes the brain activity not only during the stimulus processing stage but during the prestimulus period as well. The prestimulus *θ* and *α*-band power grow in the course of the experiment that indicates the mental fatigue caused by the increased time spent performing the task. The prestimulus *β*-band power increases in the right hemisphere that may reflect the activation of the attention network. Finally, we show that the prestimulus *β*-band power increases locally in the region, engaged in the ongoing stimulus processing, that may reflect the neuronal adaptation [19]. We hypothesize that the neuronal adaptation is enhanced in the course of the experiment reducing the cognitive demands required to activate the stimulus-related brain regions.

## Materials and methods

### Participants

Twenty healthy subjects (11 males and 9 females) aged from 26 to 35 with normal or corrected-to-normal visual acuity participated in the experiments. All of them provided written informed consent in advance. All participants were familiar with the experimental task and did not participate in similar experiments in the last six months. The experimental studies were performed under the Declaration of Helsinki and approved by the local Research Ethics Committee of the Innopolis University.

### Visual stimuli

The ambiguous visual stimulus was the Necker cube [20, 21]. A subject without any perceptual abnormalities perceives the Necker cube as a 3D-object due to the specific position of the cube’s edges. The Necker cube can be interpreted as left- or right-oriented depending on the contrast of the inner edges. The contrast of the three middle edges centered in the left middle corner was used as a control parameter *a* ∈ [0, 1]. The values *a* = 1 and *a* = 0 correspond, respectively, to 0 (black) and 255 (white) pixels’ luminance of the inner lines using the 8-bit gray-scale palette. Therefore, the control parameter was defined as *a* = *g/*255, where *g* is the brightness of the inner lines. In our experiment, we used the Necker cube images with eight different ambiguity levels (Fig. 1, A). Half of them (*a* ∈ {0.0, 0.15, 0.4, 0.45}) are considered as left-oriented (LO) and another half (*a* ∈ {0.55, 0.6, 0.85, 1}) as right-oriented (RO). While for *a* ≈ 0 and *a* ≈ 1 (low ambiguous (LA) images) the cubes can easily be interpreted as left- and right-oriented, for *a* ≈ 0.5 the identification of the cube orientation is a more difficult task since we deal with highly ambiguous (HA) images [22, 23]. The 14.2-cm Necker cubes were drawn by black and gray lines on a white background at the center of a 24” BenQ LCD monitor with a spatial resolution of 1920 × 1080 pixels and a 60-Hz refresh rate. The subjects located at a 70–80 cm distance from the monitor with a visual angle of approximately 0.25 rad.

**Fig 1.**
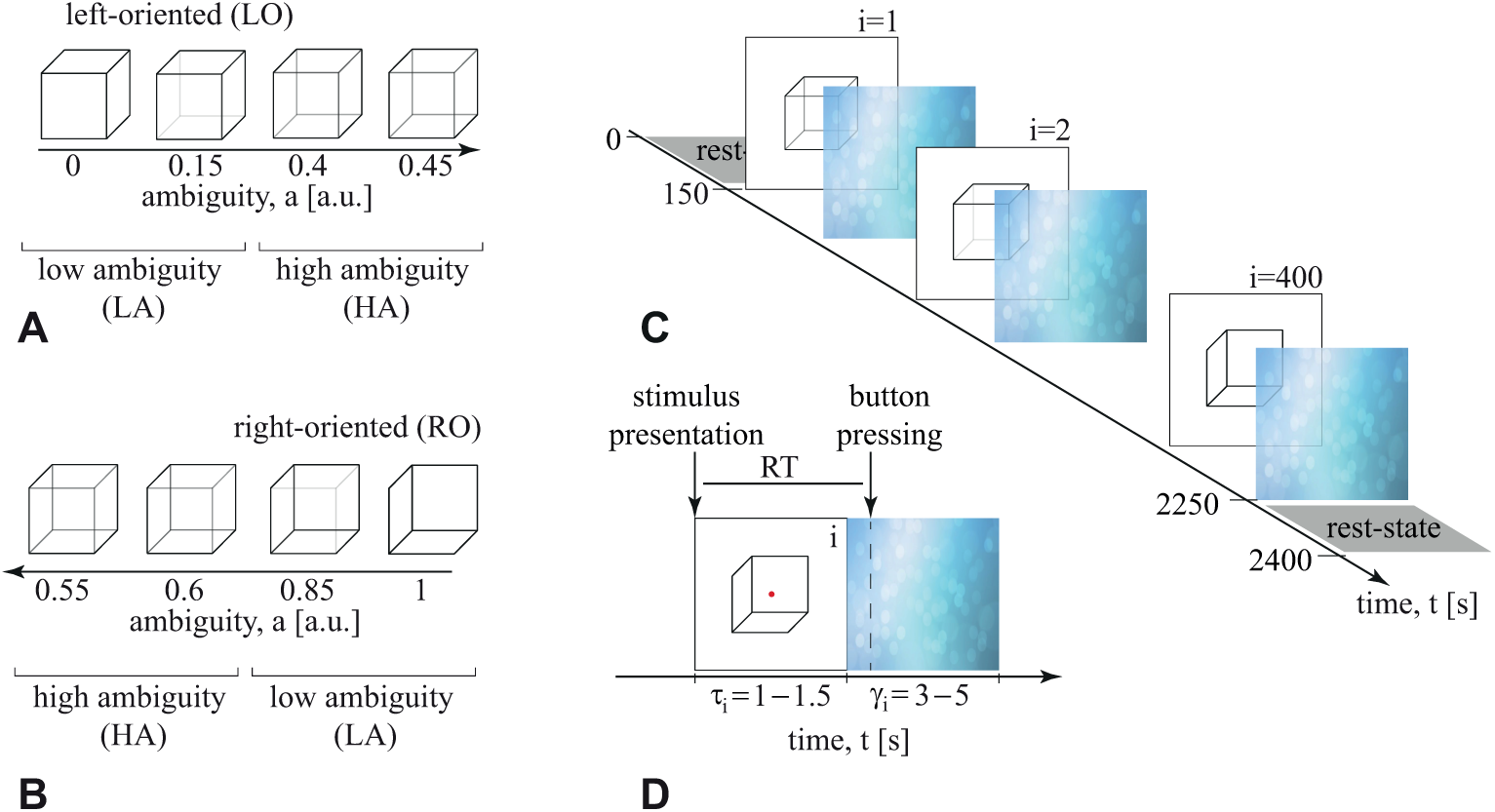
Visual stimuli. The set of visual stimuli, Necker cubes, with different degree of ambiguity *a* including high-ambiguity (HA) and low-ambiguity (LA) stimuli. A: left-oriented (LO) Necker cubes. B: right-oriented (RO) Necker cubes. C: Schematic illustration of experimental sessions. *τ*_*i*_ is the duration of the *i*-th cube presentation, *γ*_*i*_ is the interval between the *i*-th and (*i* + 1)-th presentations. D: Response time (RT) definition.

### Experimental protocol

The whole experiment lasted ≈ 40 min for each participant, including short recordings of the resting EEG state (≈ 150 s) before and after the main part of the experiment. During experimental sessions, the Necker cubes with predefined values of *a* (chosen from the set in Fig. 1, A) were randomly demonstrated 400 times, each cube with a particular ambiguity was presented about 50 times.

The scheme of the experimental session is shown in Fig. 1, B. Each *i*-th stimulus was presented for the time interval (*τ*_*i*_), followed by the time interval (*γ*_*i*_) of the abstract image presentation (see Fig. 1, C). The participants were instructed to press the left or the right key with respectively left or right hand to report their interpretation of the orientation (left or right) of each cube. The consecutively presented images affect the perception of previously demonstrated cubes. For example, if the subject observed several left-oriented cubes in a row, then his/her perception was stabilized to the left-oriented cube, even if the next cube was right-oriented. Such phenomenon is referred to as a stabilization effect [24]. To reduce this effect, the duration of the stimulus exhibition varied in the range of *τ* ∈ [1, 1.5] s. Moreover, a random variation of the control parameter *a* also prevented the perception stabilization. Lastly, to draw away the observer’s attention and make the perception of the next Necker cube independent of the previous one, different abstract pictures were exhibited for about *γ* ∈ [3, 5] s between subsequent demonstrations of the Necker cube images. For each cube, we estimated a behavioral response by measuring the response time (RT), which corresponded to the time passed from the stimulus presentation to the button pressing (Fig. 1, D) and the error (if the subject’s interpretation differs from the actual cube orientation).

### EEG acquisition and preprocessing

We recorded the EEG signals using the monopolar registration method (a 10–10 system proposed by the American Electroencephalographic Society [25]). We recorded 31 signals with two reference electrodes A1 and A2 on the earlobes and a ground electrode N just above the forehead. We used the cup adhesive Ag/AgCl electrodes placed on the “Tien–20” paste (Weaver and Company, Colorado, USA). Immediately before the experiments started, we performed all necessary procedures to increase skin conductivity and reduce its resistance using the abrasive “NuPrep” gel (Weaver and Company, Colorado, USA). Usually, the impedance varied within a 2–5 kΩ interval during the experiment. The electroencephalograph “Encephalan-EEG-19/26” (Medicom MTD company, Taganrog, Russian Federation) with multiple EEG channels and a two-button input device (keypad) performed amplification and analog-to-digital conversion of the EEG signals. This device possessed the registration certificate of the Federal Service for Supervision in Health Care No. FCP 2007/00124 of 07.11.2014 and the European Certificate CE 538571 of the British Standards Institute (BSI). The raw EEG signals were sampled at 250 Hz, filtered by a band-pass FIR filter with cut-off points at 1 Hz (HP) and 100 Hz (LP) and by a 50-Hz notch filter by embedded a hardware-software data acquisition complex. Eyes blinking and heartbeat artifact removal was performed by Independent Component Analysis (ICA) using EEGLAB software [26]. The recorded EEG signals presented in proper physical units (millivolts) were segmented into two sets of 4-s trials, including 2-s prestimulus (baseline) activity and 2-s poststimulus activity. Data were then inspected manually and corrected for remaining artifacts.

### Trials selection and experimental conditions

After the EEG preprocessing procedure, we excluded some trials due to high-amplitude artifacts. To keep the number of EEG trials constant for each cube ambiguity, we considered 320 trials out of the initial 400, including 40 trials for each ambiguity. To define the experimental conditions, we divided the whole experimental session into six non-overlapping fragments (*T*_*i*_) of the equal length. For each fragment, we selected 40 trials with an equal proportion of RO and LO stimuli, as well as LA and HA stimuli. Such a selection of the trials enables neglecting the effects caused by the cube orientation and complexity. For each segment, we averaged RT across the 20 trials corresponding to LA and HA stimuli and compared the mean RT values across the conditions *T*_1_ … *T*_6_ in the group of subjects.

### EEG time-frequency analysis

Time-frequency analysis of EEG trials was carried out using Morlet wavelet with the number of cycles for each frequency *f* as *f/*2. The wavelet power *E* was calculated in the 4 − 30 Hz frequency band and averaged over the frequency bands of interest: *θ* (4 − 8 Hz), *α* (8 − 12 Hz) and *β* (15 − 30 Hz). The prestimulus oscillatory activity was averaged over the time interval from −0.5 s to 0 s. The poststimulus oscillatory activity was contrasted by the activity in the prestimulus period subtracting the mean of baseline values followed by dividing by the mean of baseline values (‘*percent*’ mode) and considered in the time interval from 0 s to 0.5 s. The results of the time-frequency analysis were averaged over trials for each subject belonging to each fragment (*T*_1_ … *T*_6_).

Following the Ref. [27], a group-level spatio-temporal non-parametric cluster-based statistical test was used to address the statistical significance of the *θ, α* and *β* oscillatory activity changes in the prestimulus and poststimulus states between the different conditions *T*_*i*_. Here, a pairwise comparison of samples between the conditions was performed via paired sample t-test with a critical level of *p*_*pairwise*_ set to 0.01. The critical level for non-parametric cluster-based statistical test *p*_*cluster*_ was set to 0.05. The number of random permutations was 2000. The time-frequency analysis and non-parameteric cluster-based statistical tests were performed using MNE package for Python 3.7 [28]. To estimate the inter-hemispherical asymmetry of prestimulus oscillatory activity in the *θ, α* and *β* bands between the conditions *T*_*i*_, the EEG spectral power of the corresponding frequency bands was averaged over the EEG sensors placed in the left hemisphere (LH) {O1, P3, T5, CP3, TP7, C3 T3, FC3 FT7, F3, F7, FP1} and the right hemisphere (RH) {O2, P4, T6, CP4, TP8, C4, T4, FC4, FT8, F4, F8, FP2}.

## Results

### Behavioral results

We compared RT across the fragments *T*_*i*_ via a repeated-measures ANOVA with two factors: ambiguity (LA and HA) and fragment (*T*_1_ … *T*_6_). As a result, ANOVA with the Greenhouse Geisser correction revealed the significant main effect for fragment (*F*_2.31,44.05_ = 9.63, *p* < 0.001) and ambiguity (*F*_1,19_ = 59.66, *p* < 0.001), whereas the interaction effect *fragment* × *ambiguity* was insignificant (*F*_2.47,46.93_ = 1.06, *p* = 0.366). We concluded that RT changed between the fragments similarly regardless of the stimulus ambiguity. Therefore, for further analysis, we combined LA and HA stimuli on each fragment. The post-doc analysis via the Bonferroni-corrected paired-samples *t*-test revealed a monotonous decrease of RT across the fragments *T*_1_ … *T*_6_ (Fig. 2, A) with the maximal statistically significant difference achieved between the beginning (*T*_1_) and the end (*T*_6_) of the experiment (M= 0.189, SE= 0.043, *t*(19) = 4.318, *p* = 0.006). The detailed analysis of pairwise differences revealed that 18/20 subjects shown an effect in the same direction as the group (Fig. 2, B).

**Fig 2.**
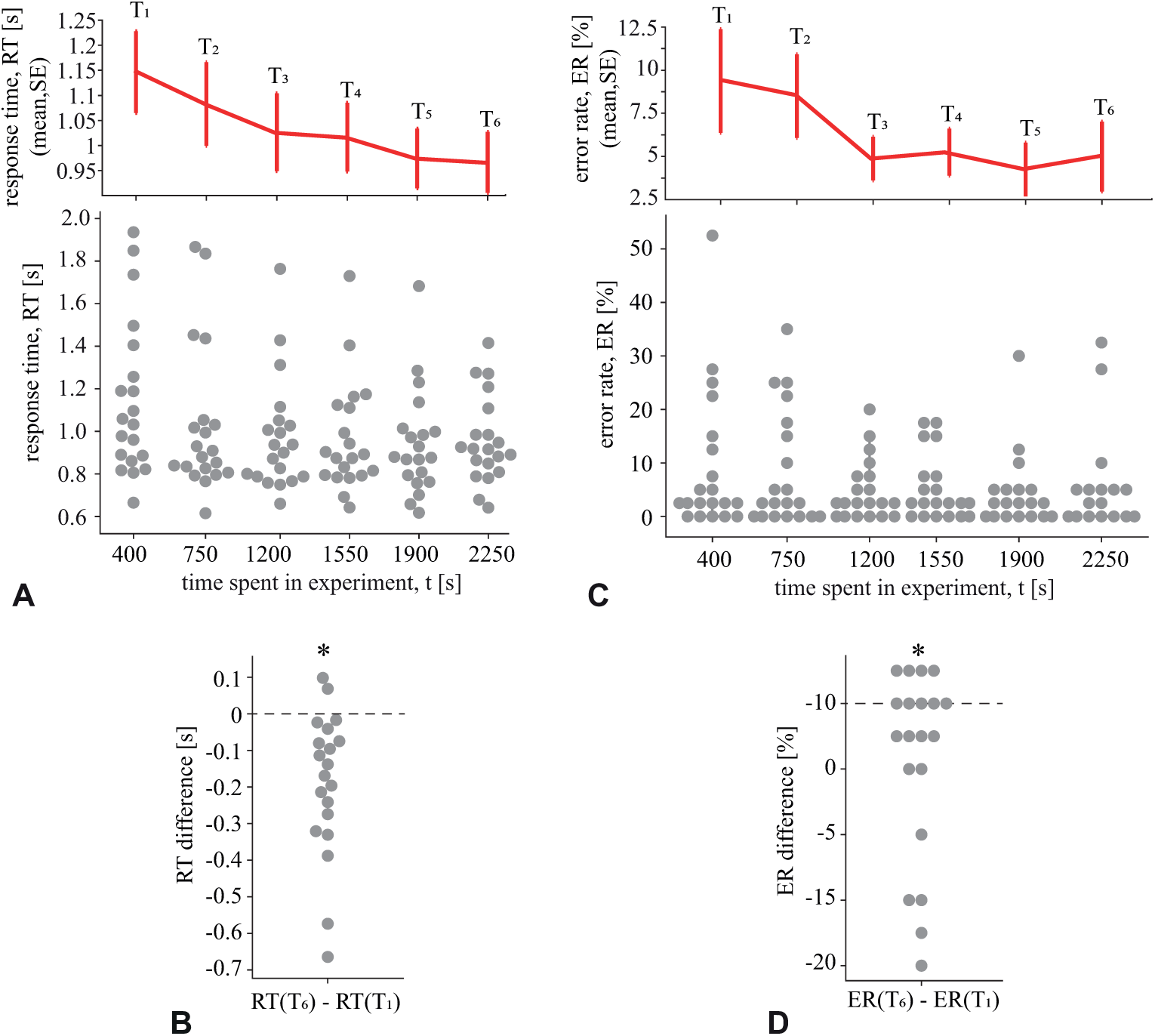
Behavioral performance in the course of the experiment. A: The response time (RT) for each fragment (*T*_1_ … *T*_6_) of the experimental session. The upper panel demonstrates the group mean±SE, while the lower panel reflects the data of all subjects. B: The error rate (ER) for each fragment of the experimental session (mean±SE and the individual values). C: The change of RT between the last (*T*_6_) and the first (*T*_1_) fragments. D: The change of ER between these fragments.

A similar tendency was observed for the error rate (ER) (Fig. 2, C). ANOVA with the Greenhouse Geisser correction revealed a significant effect for the fragment (*F*_2.75,52.38_ = 3.386, *p* = 0.028). The posthoc analysis with the paired samples *t*-test displayed the maximal statistically significant difference between *T*_1_ and *T*_6_ (M= 1.74, SE= 0.64, *t*(19) = 2.733, *p* = 0.013). The Bonferroni-corrected *p*-value was 0.198. The distribution of pairwise differences reflected that 11/20 subjects followed the group tendency and 5/20 subjects shown no effect (Fig. 2, D).

The obtained results demonstrated that both the RT and the ER decreased in the course of the experiment. We supposed that the most significant difference in the neuronal activity should be observed between the fragments *T*_1_ and *T*_6_. For these conditions, we compared RT between the LO and RO cubes. The repeated-measures ANOVA with the Greenhouse Geisser correction revealed a significant change in RT between *T*_1_ and *T*_6_ (*F*_1,19_ = 20.714, *p* < 0.001). In contrast, RT changed insignificantly between LO and RO cubes (*F*_1,19_ = 1.56, *p* = 0.227). Finally, we found that the interaction effect was also insignificant (*F*_1,19_ = 0.084, *p* = 0.775).

This analysis evidenced that a change in RT between the *T*_1_ and *T*_6_ conditions was independent of the cube ambiguity and orientation. Given above, cortical activity was compared between *T*_1_ and *T*_6_ conditions based on 40 trials per condition, including the equal proportion of the LA and HA stimuli as well as LO and RO stimuli.

### Rest-state and prestimulus activity

First, we analyzed how the subject’s condition changed in the course of the experiment. The human condition affects brain activity regardless of the presented stimuli or the task. Therefore, we analyzed the EEG spectral power in three conditions: rest-state (the phase of the experiment before the first stimulus was presented), prestimulus state at the beginning of the experiment (*T*_1_) and prestimulus state at the end of the experiment (*T*_6_). For each condition, we considered the set of 40 EEG trials (0.5-s length). Fig. 3 illustrates the EEG spectral power in the *θ, α*, and *β* bands in the left (LH) and right hemispheres (RH) for these conditions.

**Fig 3.**
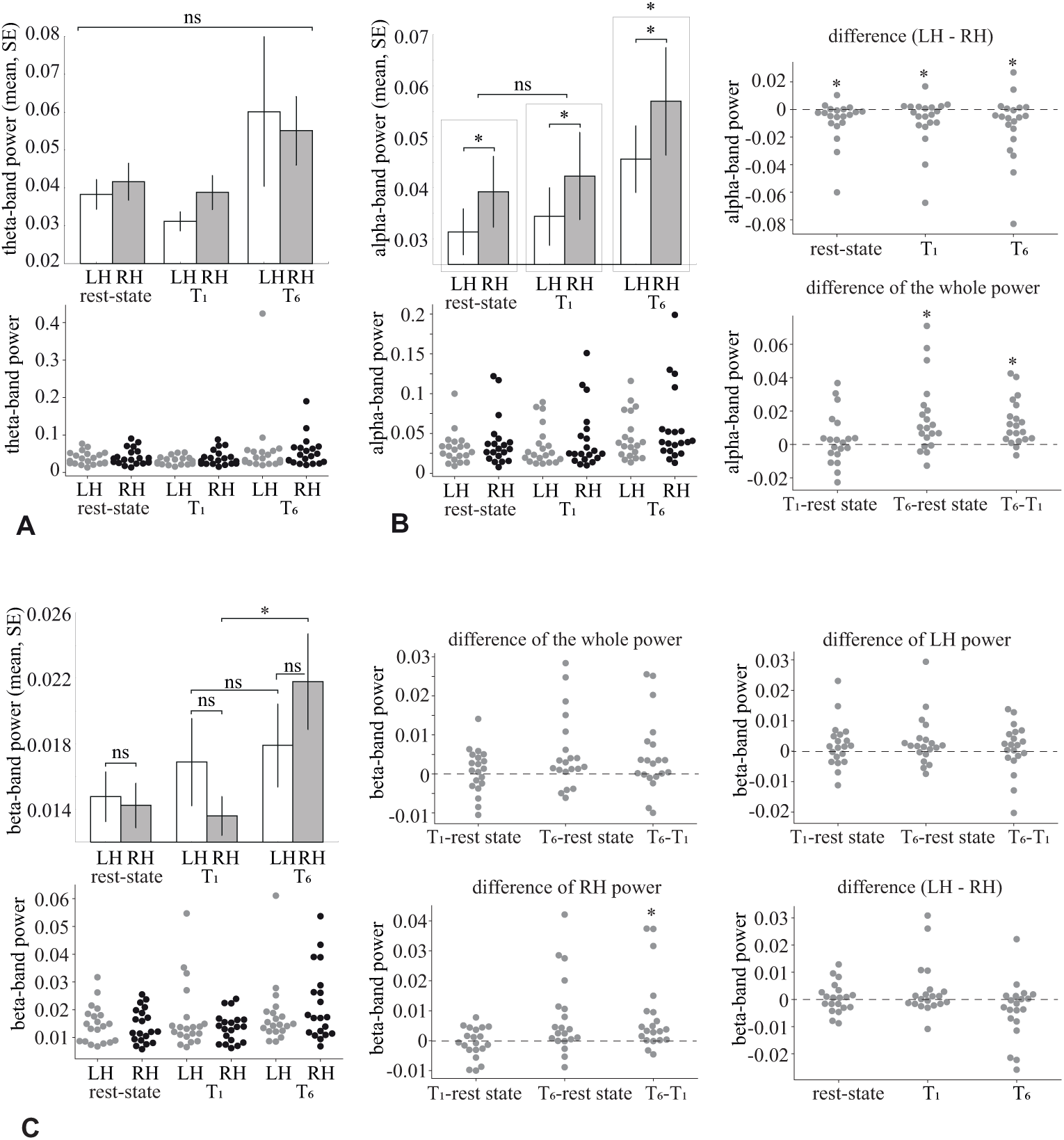
Rest-state and prestimulus wavelet power. A: *θ*-band power in the left hemisphere (LH) and the right hemisphere (RH) during three conditions (rest-state, prestimulus period at the beginning (*T*_1_) and prestimulus period at the end (*T*_6_) of the experiment). The upper panel represents group mean±SE, and the lower panel shows the data of all subjects. B: *α*-band power across the same conditions (mean±SE and the individual values). Differences between the LH and RH *α*-band power for the rest-state, *T*_1_ and *T*_6_ conditions. The differences of the *α*-band power in both hemispheres between the different conditions. C: *β*-band power in the left hemisphere (LH) and the right hemisphere (RH) for the rest-state, *T*_1_ and *T*_6_ conditions (mean±SE and the individual values). The differences of the *β*-band power in both hemispheres between the different conditions. The differences of the RH *β*-band power between the different conditions. The differences of the LH *β*-band power between the different conditions. Differences between the LH and RH *β*-band power for the rest-state, *T*_1_ and *T*_6_ conditions.

In the *θ*-band (Fig. 3, A) the repeated measures ANOVA with the Greenhouse-Geisser correction revealed insignificant main effect for the hemisphery (*F*_1.00,19.00_ = 0.18, *p* = 0.676) and for the condition (*F*_1.058,20.096_ = 0.18, *p* = 0.124). The interaction effect hemisphere×condition was also insignificant (*F*_1.067,20.268_ = 0.823, *p* = 0.383).

In the *α*-band (Fig. 3, B), the repeated measures ANOVA with the Greenhouse-Geisser correction revealed a significant main effect for the condition (*F*_1.46,27.84_ = 9.713, *p* = 0.002) and the hemisphere (*F*_1.00,19.00_ = 4.944, *p* = 0.039). The interaction effect condition×hemisphere was insignificant (*F*_1.27,24.146_ = 1.378, *p* = 0.264). The post-hoc analysis with Bonferroni-corrected paired samples *t*-test demonstrated that *E*_*α*_ (RH) exceeded *E*_*α*_ (LH) (M= 9.06 × 10^3^, SE= 4.07 × 10^3^, *p* = 0.039). The distribution of the pairwize differences (Fig. 3, B) evidenced that 14/20 subjects followed the group tendency at the rest-state, while 12/20 and 14/20 subjects followed the group tendency at *T*_1_ and *T*_6_ conditions, respectively.

*E*_*α*_ in the rest-state did not differ significantly from *E*_*α*_ (*T*_1_) (*p* = 1.0), while *E*_*α*_ (*T*_6_) significantly exceeded both *E*_*α*_ (*T*_1_) (M= 15.9 × 10^3^, SE= 4.84 × 10^3^, *p* = 0.012) and *E*_*α*_ during the rest-state (M= 12.8 × 10^3^, SE= 3.01 × 10^3^, *p* = 0.001). The pairwize diffeences evidenced that 15/20 and 18/12 subjects followed the group effect when *T*_6_ was compared to the rest-state and *T*_1_ conditions.

We concluded that participants exhibited the right-lateralization of the *α*-band power during the rest-state and in the prestimulus state during the experiment. The overall prestimulus *α*-band power at the beginning of the experiment did not differ from the rest-state. The overall *α*-band power in the prestimulus period increased in the course of the experiment.

In the *β*-band (Fig. 3, C), repeated measures ANOVA with Greenhouse-Geisser correction revealed a significant main effect for the condition (*F*_1.531,29.094_ = 4.654, *p* = 0.025), and insignificant main effect for the hemisphere (*F*_1.00,19.00_ = 0.0, *p* = 0.988). The interaction effect condition×hemisphere was also significant (*F*_2,38_ = 7.619, *p* = 0.002). The post-hoc analysis with Bonferroni-corrected paired samples *t*-test demonstrated that *E*_*β*_ in the rest-state did not differ significantly from *E*_*β*_ (*T*_1_) (*p* = 1.0), and *E*_*β*_ (*T*_6_) did not significantly differ from both *E*_*β*_ (*T*_1_) (*p* = 0.136) and *E*_*β*_ during the rest-state (*p* = 0.066). At the same time, analysis of the pairwize differences (Fig. 3, C) revealed that 16/20 and 18/20 subjects exhibited higher *E*_*β*_ (*T*_6_) when compared to the *E*_*β*_ at the rest-state and to the *E*_*β*_ (*T*_1_), respectively.

The paired samples *t*-test revealed that *E*_*β*_ (LH) and *E*_*β*_ (RH) differed neither for the rest-state (*t*(19) = 0.423, *p* = 0.677) nor for the prestimulus period at the beginning (*t*(19) = 1.49, *p* = 0.151) and at the end (*t*(19) = −1.62, *p* = 0.121) of the experiment.

A paired-samples *t*-test was conducted to compare the *β*-band power in the right hemisphere *E*_*β*_ (RH) across the conditions. We revealed that *E*_*β*_ (RH) at *T*_1_ did not differ from the *E*_*β*_ (RH) at the rest-state (*t*(19) = 0.562, *p* = 0.58). *E*_*β*_ (RH) at *T*_6_ significantly exceeded *E*_*β*_ (RH) at *T*_1_ (M= 8.17 × 10^3^, SE= 2.81 × 10^3^, *t*(19) = 2.899, *p* = 0.009). Analysis of the pairwize differences (Fig. 3, C) revealed that 18/20 subjects shown this effect. The difference of *E*_*β*_ (RH) between *T*_6_ and rest-state was also significant (M= 7.53 × 10^3^, SE= 2.85 × 10^3^, *t*(19) = 2.639, *p* = 0.016). This effect was observed for 17/20 subjects.

A paired-samples *t*-test was conducted to compare the *β*-band power in the left hemisphere, *E*_*β*_ (LH), in the rest-state, *T*_1_ and *T*_6_ conditions. We revealed that *E*_*β*_ (LH) at *T*_1_ did not differ from the *E*_*β*_ (LH) at the rest-state (*t*(19) = 1.273, *p* = 0.218). *E*_*β*_ (LH) at *T*_6_ did not differ from both *E*_*β*_ (LH) at *T*_1_ (*t*(19) = 0.565, *p* = 0.579) and at the rest-state (*t*(19) = 1.733, *p* = 0.099).

Finally, according to the paired-samples *t*-test, *E*_*β*_ (RH) did not significantly differ from *E*_*β*_ (LH) at the rest-state (*t*(19) = −0.423, *p* = 0.677) as well as at *T*_1_ (*t*(19) = −1.495, *p* = 0.151) and *T*_6_ (*t*(19) = 1.624, *p* = 0.121) conditions.

The obtained results demonstrated that the *β*-band power did not change at the beginning of the experiment when compared to the rest-state. In the course of the experiment, we observed an increase in the prestimulus *β*-band spectral power in the right hemisphere.

Fig. 4 shows the topograms of *t*-values reflecting the power changes in the *θ*-, *α*−, and *β*-frequency bands in the *T*_6_ versus *T*_1_ conditions.

**Fig 4.**
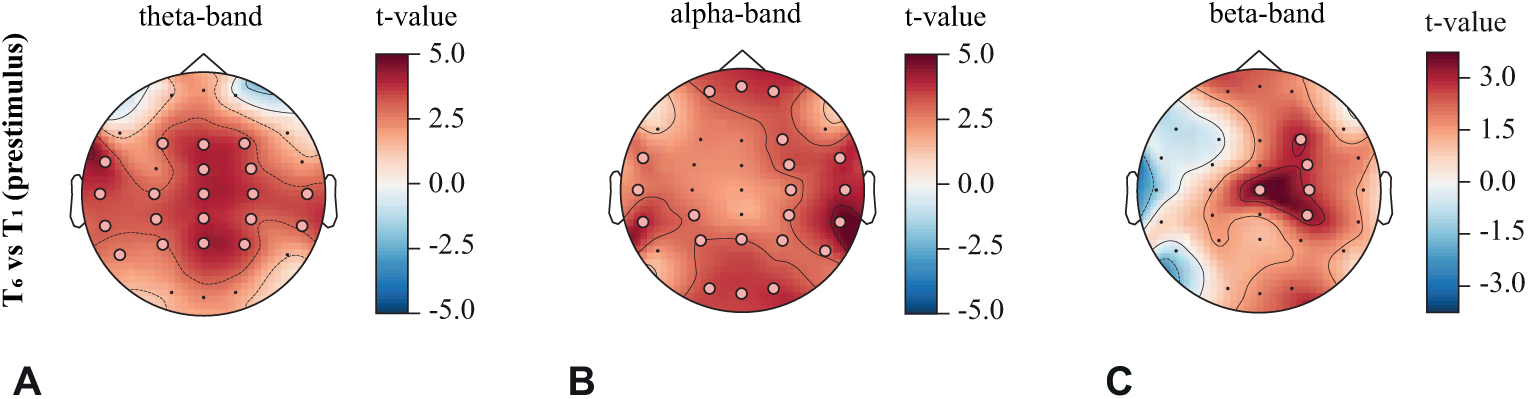
Prestimulus wavelet power distribution. The *t*-value and the EEG channel clusters representing significant changes in (A) *θ*-, (B) *α*- and (C) *β*-frequency band spectral power at the end (*T*_6_) versus the beginning (*T*_1_) of the experiment (*p*_*pairwise*_ < 0.01, *p*_*cluster*_ < 0.05)

In the *θ*-band (Fig. 4, A), the cluster-based statistical analysis with permutations revealed that the prestimulus state at the end of the experiment is characterized by the increasing power over the EEG channel cluster in the parietal, sensorimotor, frontal and temporal areas.

In the *α*-band (Fig. 4, B), a significant increase of *α*-band power at the end of the experiment was observed for the channel clusters in the occipito-parietal area, frontal area, and temporal areas.

In the *β*-band (Fig. 4, C), the prestimulus state at the end of the experiment was characterized by a significant (*p* < 0.01) increase in the *β*-band spectral power for the right-lateralized channel cluster including channels F4, FC4, C4, CP4, Cz.

### Task-related activity

The task-related cortical activity on the EEG sensor level was analyzed by comparing the topograms of the *θ*-, *α*- and *β*-band spectral power between the beginning (*T*_1_) and the end (*T*_6_) of the experiment. The Fig. 5 illustrates the channels clusters (circles) and the corresponding *t*-values (color scale) obtained during the 0.5-s interval following the presentation of the visual stimulus. Each topogram represented the spectral power averaged over a 0.1-s window mentioned in the legend under the topograms.

**Fig 5.**
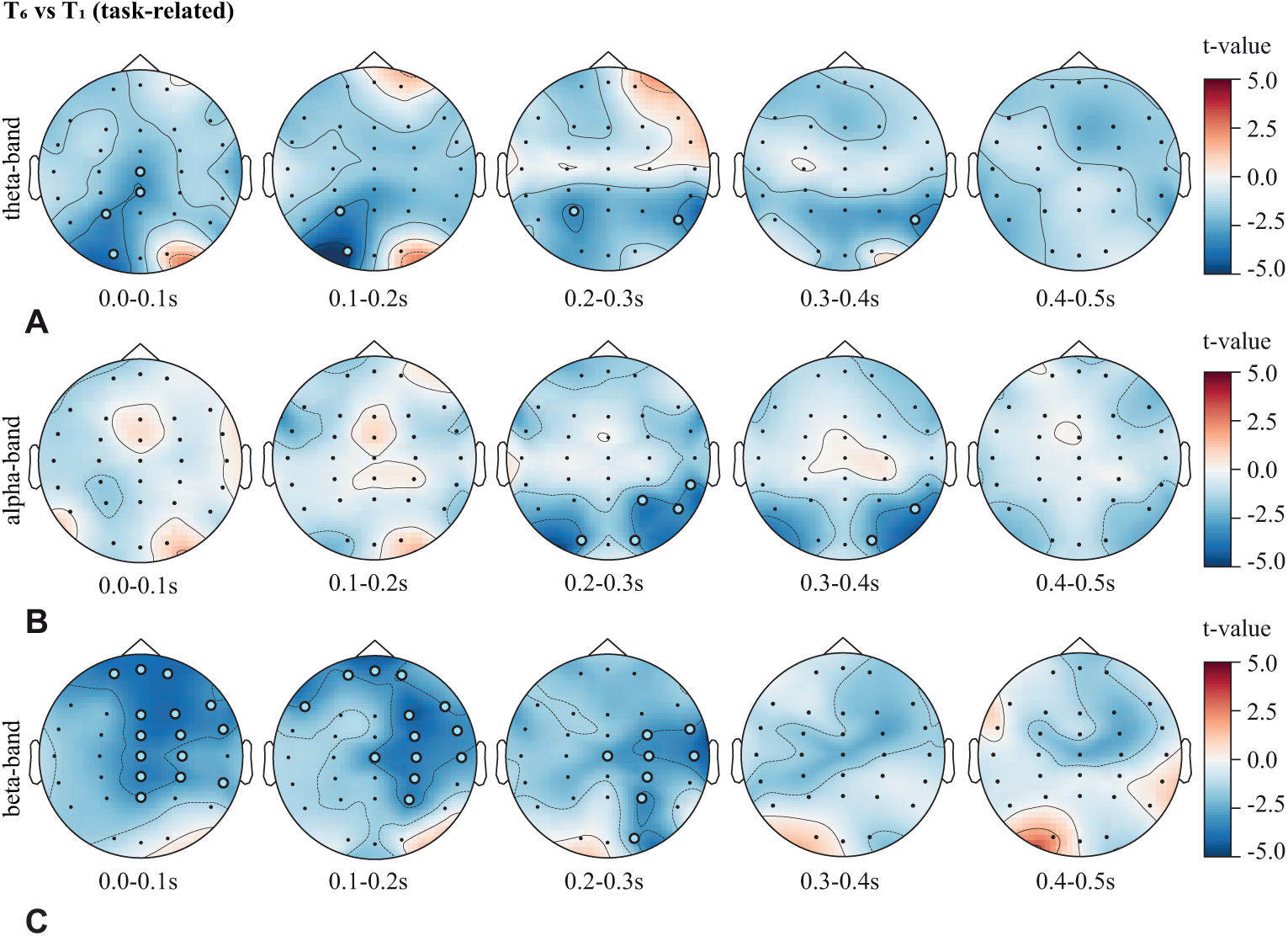
Task-related spatio-temporal activity. The *t*-value and the EEG channel clusters representing significant changes in the wavelet power in the (A) *θ*-, (B) *α*- and (C) *β*-frequency bands at the end (*T*_6_) versus the beginning (*T*_1_) of the experiment (*p*_*pairwise*_ < 0.01, *p*_*cluster*_ < 0.05)

One could see that the clusters had different spatio-temporal properties in the different frequency bands. At the same time, no positive clusters appeared in these bands, indicating the overall increase of the task-related neuronal activation at the beginning of the experiment.

In the *θ*-band (Fig. 5, a), the earlier processing stage (*t* < 0.1 s) was characterized by the left-lateralized occipito-parietal cluster (O1 and P3 channels) and the mid-line centro-parietal cluster (CPz and Cz channels). For (0.1 < *t* < 0.2 s), the only left-lateralized occipito-parietal (O1 and P3) cluster was observed. The further processing (*t* > 0.2 s) was associated with the increased *t*-value over the parietal and temporal EEG sensors. For (0.2 < *t* < 0.3 s), there were two clusters (P3 and P8), while for (0.3 < *t* < 0.4 s), the only one cluster (P8) remained. Finally, for (*t* > 0.4 s), no clusters were found in the *θ*-frequency band.

The *α*-band power (Fig. 5, b) remained unchanged during the first 0.2 sec after the stimulus presentation. Then, for 0.2 − 0.3 s, we observed the bilateral negative cluster in the occipital area, including O1 and O2 EEG channels and the right-lateralized parieto-temporal cluster, including P4, P8, and TP8 EEG channels. For the 0.3 − 0.4 s, the size of the observed cluster decreased, and it became right-lateralized. Finally, the further increase of the processing time led to the cluster disappearance. The obtained results evidenced that at the end of the experiment, the task-related cortical activity was characterized by the increased *α*-band power in the occipital and right temporal areas during 0.2 − 0.3 s after the stimulus presentation.

In the *β*-frequency band (Fig. 5, c), the negative cluster was observed for the first 0.3 s after the visual stimulus presentation. At the earlier sensory processing stage (*t* < 0.1 s) it included the bilateral prefrontal cortical channels (Fp1, Fpz, Fp2), the right-lateralized frontoparietal area (Fz, F4, F8, FCz, FC4, Cz, C4, CPz, CP4, Pz) and the right temporal area (FT8, TP8). During the 0.1 − 0.2 s, the observed cluster was bilateral in the prefrontal cortex, but right-lateralized in the temporal region. For 0.2 − 0.3 s, the frontal cluster disappeared, and one could observe the remaining right-lateralized cluster in the occipital, parietal, and temporal cortex. Finally, no clusters were found for *t* > 0.3 s in the *β*-frequency band.

## Discussion

Our results demonstrate that during a prolonged visual stimuli classification task, subjects enhance their performance in terms of the reduced reaction time and decision accuracy. This observation is not trivial since a prolonged cognitive activity may induce mental fatigue and, therefore, cognitive decline. Thus, we suppose that the brain implements a strategy for the optimal utilization of cognitive resources to resist mental fatigue and increase behavioral performance.

At the beginning of the experiment, we observe enhanced spectral power in the *θ, α*, and *β* frequency bands during the stimulus processing. The high *β* power has been observed in the right temporoparietal cortex and bilaterally in the prefrontal cortex, while high *α* and *θ* power – in the occipital and in both occipital and parietal areas, respectively.

The neuronal activity in the temporoparietal regions subserves the processing and storage of visuospatial information [29, 30]. At the same time, the activation of the prefrontal cortex indicates that the task accomplishing requires additional cognitive resources. It is also confirmed by an increase of the task-related *θ*-band activity [31]. The task-related occipital *α*-power is associated with selective attention during stimulus processing. The low *α*-band power reflects the effective processing of attended stimuli [32]. Having summarized, we suppose that at the beginning of the experiment, stimulus processing engages the resources of the frontoparietal cortical network. Moreover, the lower task-related *α*-band power in the occipital area at the end of the experiment is possible to reflect the enhanced brain ability to respond to the attended stimulus effectively.

The prestimulus activity in the *θ, α*, and *β*-bands changes in the course of the experiment. The prestimulus *α*-band power is higher in the right hemisphere during the rest-state and the prestimulus period. This lateralization doest not change in the course of the experiment. At the same time, *α*-band power grows during the experiment over the majority of the EEG sensors in both hemispheres. The *θ*- and *β*-band activity is bilateral in the rest-state and prestimulus period. However, at the end of the experiment, the *β*-band power increases in the right hemisphere, while the *θ*-band power grows bilaterally in the frontal-central, parietal, and temporal regions.

The prestimulus activity in the *α* and *β* bands is usually analyzed to characterize the subject’s attention. A wide body of literature shows that *α* and *β*-band activities are relevant to attention in general and not restricted to the visual stimuli processing [6, 33–35]. Attention modulates the prestimulus *α*- and *β*-band power [6, 33, 36] and affects the stimulus processing accuracy. Thus, low *α*- and high *β*-band power during the prestimulus period is beneficial for sensory perception [5, 6]. The prestimulus *θ*- and *α*-band activities can serve as the markers of mental fatigue, i.e., the increased power in these bands reflects the subject’s mental fatigue and causes the performance decrement [3]. Taken together, the low prestimulus *θ*- and *α*-band power and the high *β*-power should predict the high performance and vice versa. On the contrary, we report that the high prestimulus spectral power in the *θ*-, *α*- and *β*-frequency bands accompany the performance increment.

According to [37], the lateralized *α*-band power reflects the attentional breadth during the stimulus processing. The attentional breadth characterizes the subject’s ability to focus attention either on the global level (e.g., the forest) or local elements that make up the stimulus (e.g., the trees) [38]. Recent studies demonstrate that greater local attentional bias is observed after activating the left hemisphere, and a greater global bias is observed after activating the right hemisphere [39]. The *α*-band power over the right and left frontal-central sensors correlate with attentional breadth. The greater left frontal *α* power corresponds to faster responses to local targets and greater right frontal *α* power – to faster responses to global targets [40]. Previous findings suggest that prestimulus and rest-state *α* power lateralization [41] also predict the subject’s performance in a global-local attention task [37].

We report the high *α*-band activity in the right hemisphere at rest as well as in the prestimulus state. Moreover, the lateralization does not change in the course of the experiment. Therefore, we can conclude that the subjects which initially have right-lateralized alpha-band power exhibit the global attentional bias and do not change it during the experiment. It coincides with the existing literature claiming that the attentional bias remains stable for at least ten days in multiple global/local tasks [42].

The increased prestimulus *α* power in the course of the experiment can be an electrophysiological indicator of mental fatigue [43]. The prolonged task induces mental fatigue, increasing reaction times, misses, and false alarms. As discussed above, high *α*-band power predicts the low performance of the ongoing stimulus processing. Thus, it can be supposed, that increased *α*-band power at the end of the experiment is associated with mental fatigue, which should lead to the performance decline. The increased *θ*-band power, together with *α*-band power, can also manifest the mental fatigue. According to [3], the prestimulus *θ*-band power in the frontal area grows in the course of the experiment and negatively correlates with the behavioral performance. Another paper [44], relates mental fatigue with the increased *θ*-band power in the frontal-central and parietal regions.

High prestimulus *β*-band power in the right hemisphere can be related to the increasing human attention. There is a view that the human attentional brain network is overall lateralized to the right hemisphere [45]. Some components of the attention (e.g., alerting and disengaging functions) are bilateral [46], while others (e.g., orienting and executive functions) are biased to the right hemisphere [47]. According to [48], the executive functions subserve an interplay between alerting and orienting functions to maintain the state of readiness and focus attention towards the relevant features of a stimulus.

Taken together, the increased orienting and executive components of attention in the course of the experiment can be the possible reasons for the increase in behavioral performance. In this context, the performance can grow due to the enhanced effectiveness of the relevant stimulus features selection. At the same time, high prestimulus *θ*- and *α*-power contradict the definition of attentional state in general. Therefore, we cannot conclude unequivocally that the brain increases its attentional properties in the course of the experiment.

Finally, we introduce one more possible explanation for the increased performance in the course of the experiment. We observe that the prestimulus *β*-band power at the end of the experiment increases locally in the region, which is more engaged during the stimulus processing. Hence, we suppose that the preactivation of the stimulus-related cortical areas reduces the cognitive demands required for their activation during the stimulus processing ([49]). The preactivation of the stimulus-related neuronal ensembles before the stimulus processing is also known as the neuronal adaptation (NA). The NA is considered as the evidence for the predictive coding theory [50]. NA is observed when the same visual stimulus is repeatedly presented with a brief interval and causes the reduced neural response to repeated compared with unrepeated stimuli [19]. The NA is supposed to arise from at least two types of neural activity. One explanation is that only the part of the neuronal ensemble is sensitive to stimulus recognition. Thus, the neurons that are not critical for recognizing the stimulus decrease their responses as the stimulus is repeatedly presented. In contrast, the neuronal populations carrying essential information continue to give a robust response. As a result, the mean firing rate becomes attenuated by stimulus repetition. This theory is supported by the first studies of unit cell recordings [51]. The alternative explanation is the stimulus repetition reduces the response in the temporal domain [19]. In this theory, the neural processing network settles to a stable response more quickly in response to a repeated than novel stimulus, because the network connections involved in producing the response have been reinforced by a previous presentation of the same stimulus [52]. The NA affects the neuronal response in the occipital [53], parietal [54], and frontal [55] cortical populations in the single-unit data as well as on the sensor level.

Taking together, we suppose that the training effect in the course of the experiment results in the preactivation of the stimulus-related regions in the *β* frequency band. The very recent work [56] also relates the *β*-band spectral power to the training. The authors report on the enhanced *β*-band power over the central, temporal and the parietal sensors during an 80–400 ms window of the poststimulus onset (post-versus pre-training trials), while we observe that the *β*-band spectral power decreases in the same areas during the same time window. At first glance, this issue can be addressed to different experimental paradigms. For example, [56] compared two experiments carried out in different days, whereas in this work, we analyze a single experimental session performed in one day. It follows that the different mechanisms stay behind the training effect in these two paradigms. Besides, different stimuli are used, i.e., we consider visual stimuli only, while [56] uses visual stimuli together with auditory stimuli. Moreover, similar to our findings [56] report on a significant decrease in the *β*-band power (post-training vs. pre-training trials) at certain conditions. The authors relate this effect to the participant’s perceptual template formation after the training and consider these results as the evidence for the predictive coding model. The predictive coding theory suggests that the brain creates sensory templates [57], which causes increasing *β*-band activity before the occurrence of an expected event that potentially reflects the mobilization of neuronal populations to encode the expected sensory inputs [58, 59].

## Supporting information

**S1 Data. EEG and behavioral data.** This structure contains 20 cells representing data of 20 subjects. Each cell includes field *Trial*, which contains 80 EEG trials recorded for 31 channels. The field *Label* contains the names of the channels. The first 40 trials are chosen from the beginning of the experiment and belong to the condition *T*_1_. The rest 40 trials are chosen from the end of the experiment and belong to the condition *T*_6_. Each trial has 4 s length, and the stimulus is presented in the middle of the trial. The field *PresentationMoment* contains the moments of the stimuli presentations and *Ambiguity* contains the stimulus ambiguity value. The field *ReactionTime* contains the reaction time for each trial, and the field *Button* contains the identification of the button pressed (1-for the left and 2-for the right). The field *Background* contains the rest-state EEG signals recorded before the stimuli presentation. EEG signals are sampled at 250 Hz, filtered by a band-pass FIR filter with cut-off points at 1 Hz (HP) and 100 Hz (LP) and by a 50-Hz notch filter. Eyes blinking and heartbeat artifact are removed with Independent Component Analysis (ICA)

## Acknowledgments

This work has been supported by the Russian Foundation for Basic Research (Grant 19-32-60042) in the part of experimental work and behavioral data analysis. AK is supported by the President Program (MK-1760.2020.2) in the part of experimental data preprocessing.

